# Species-specific molecular barriers to SARS-CoV-2 replication in bat cells

**DOI:** 10.1101/2021.05.31.446374

**Authors:** Sophie-Marie Aicher, Felix Streicher, Maxime Chazal, Delphine Planas, Dongsheng Luo, Julian Buchrieser, Monika Nemcova, Veronika Seidlova, Jan Zukal, Jordi Serra-Cobo, Dominique Pontier, Olivier Schwartz, Jiri Pikula, Laurent Dacheux, Nolwenn Jouvenet

## Abstract

Bats are natural reservoirs of numerous coronaviruses, including the potential ancestor of SARS-CoV-2. Knowledge concerning the interaction between coronaviruses and bat cells is sparse. We investigated the susceptibility of primary cells from *Rhinolophus ferrumequinum* and *Myotis* species, as well as of established and novel cell lines from *Myotis myotis*, *Eptesicus serotinus*, *Tadarida brasiliensis* and *Nyctalus noctula*, to SARS-CoV-2 infection. None of these cells were sensitive to infection, not even the ones expressing detectable levels of angiotensin-converting enzyme 2 (ACE2), which serves as the viral receptor in many mammalian species. The resistance to infection was overcome by expression of human ACE2 (hACE2) in three cell lines, suggesting that restriction to viral replication was due to a low expression of bat ACE2 (bACE2) or absence of bACE2 binding in these cells. Infectious virions were produced but not released from hACE2-transduced *M. myotis* brain cells. *E. serotinus* brain cells and *M. myotis* nasal epithelial cells expressing hACE2 efficiently controlled viral replication. This ability to control viral replication correlated with a potent interferon response. Our data highlight the existence of species-specific molecular barriers to viral replication in bat cells. These novel chiropteran cellular models are valuable tools to investigate the evolutionary relationships between bats and coronaviruses.

**Author summary:** Bats host ancestors of several viruses that cause serious disease in humans, as illustrated by the on-going SARS-CoV-2 pandemic. Progress in investigating bat-virus interactions have been hampered by a limited number of bat cell lines. We have generated primary cells and cell lines from several bat species that are relevant for coronavirus research. The varying susceptibilities of the cells to SARS-CoV-2 infection offered the opportunity to uncover some species-specific molecular restrictions to viral replication. All bat cells exhibited a potent entry-dependent restriction. Once this block was overcome by over-expression of human ACE2, which serves at the viral receptor, two bat cell lines controlled well viral replication, which correlated with the inability of the virus to counteract antiviral responses. Other cells potently inhibited viral release. Our novel bat cellular models contribute to a better understanding of the molecular interplays between bats and viruses.

## Introduction

Bats are natural hosts of numerous coronaviruses, including members of the *Betacoronavirus* genus, which comprises viruses belonging to the severe acute respiratory syndrome coronavirus (SARS-CoV) 1 and 2 lineages [1,2]. The RaTG13 virus, which shares 96.1% nucleotide sequence with SARS-CoV-2 [3], was sampled from faeces of *Rhinolophus affinis* in the Yunnan province of China in 2013[4]. RmYN02 virus, which also belongs to the RaTG13/SARS-CoV-2 lineage, was recently identified in *Rhinolophus malayanus* collected in China [1]. Other viruses belonging to this lineage have been recently identified in *Rhinolophus* bats sampled in Thailand [5] and in Cambodia [6]. SARS-CoV-2 related coronaviruses (SC2r-CoVs) are thus probably widely distributed in South-East Asia. In addition, numerous other bat species worldwide are infected with betacoronaviruses, including species of the *Myotis*, *Nyctalus*, *Tadarida* and *Eptesicus* genera [7–12].

The risk of spillback transmission of SARS-CoV-2 from humans to domestic animals or wildlife remains a major concern, as this reverse zoonotic transmission has been already documented in pet animals, tigers and gorillas in zoos, and farmed minks [13,14]. Given the likely bat origin of SARS-CoV-2, bats could be putatively at risk of spillback transmission [15]. The establishment of novel bat reservoirs would have a severe impact on wild-life conservation and public health measures.

Betacoronaviruses circulating in bats and humans use the surface receptor angiotensin-converting enzyme 2 (ACE2) to enter cells [4,16–18]. Viral binding to ACE2 is followed by the proteolytic cleavage of the viral spike (S) proteins by either the plasma-membrane resident transmembrane protease serine 2 (TMPRSS2) or the endosomal cathepsin L (CTSL)[19]. This cleavage is mandatory for the fusion between the viral and cellular membranes. Thus, localization and expression of TMPRSS2 and CTSL dictate whether the virus enters cells by fusing at the cell surface or in endosomes [19].

Several approaches have been used to predict the ability of ACE2 from phylogenetically diverse bat species to promote viral entry. First, comparison of ACE2 protein sequences from 37 bat species, including species of the genus *Rhinolophus*, predicted a low or very low ability to interact with viral S proteins[20]. Second, expressing ACE2 from dozen bat species in non-permissive mammalian cells using genuine viruses or pseudo-viruses carrying SARS-CoV-2 S proteins revealed that ACE2 from *Rhinolophus, Myotis* and *Eptesicus* species allowed viral entry [21–24], albeit often less efficiently than human ACE2. However, these approaches using *in silico* analysis or ectopic expression of bat ACE2 in human or hamster cells do not allow to draw conclusions as to which bat species might support SARS-CoV-2 replication. Other factors unique to bat cells may potentially modulate viral entry and replication. Indeed, experiments performed with cells derived from lung tissue of *Rhinolophus alcyone* and *Myotis daubentonii* showed that they were not susceptible to infection with vesicular stomatitis viruses (VSV) bearing SARS-CoV-2 S proteins [17]. Cells originating from lung and kidney tissue of *Rhinolophus sinicus* and *Eptesicus fuscus* were not permissive to SARS-CoV-2 either [25,26]. These studies underline the limitation of predicting the ability of S proteins to bind ACE2 orthologs based on computational models or ectopic expression.

Only a handful of models are available to study the replication of betacoronaviruses in bat cells. Viral replication was detected in *Rhinolophus sinicus* lung and brain cells, as well as in *Pipistrellus abramus* kidney cells [27], but viral titers were very low. By contrast, SARS-CoV-2 replicated efficiently in *R. sinicus* intestinal organoids [28], confirming further the susceptibility of *Rhinolophus* cells to the virus. Intranasal inoculation of SARS-CoV-2 in *Rousettus aegypticus* resulted in transient infection of their respiratory tract and oral shedding of the virus [29], indicating that bats unrelated to the *Rhinolophus* genus are also susceptible to the virus. Since the manipulation of bat organoid and animal models remains challenging, there is a need to develop cell lines from various organs and species to gain deeper insights into bat-virus co-evolution [30]. Here, we developed novel cellular models derived from bat species circulating widely in Europe and Asia. The varying susceptibilities of the cells to SARS-CoV-2 infection offered the opportunity to uncover some species-specific molecular restrictions to viral replication.

## Results

### Resistance to SARS-CoV-2 infection in selected bat cell lines

Species belonging to the *Rhinolophus* genus, including *R. ferrumequinum*, are known natural hosts for numerous SARS-CoV-related betacoronaviruses [9,31]. Alphacoronaviruses [10,32,33], and possibly betacoronaviruses [8], circulate in species belonging to the *Myotis* genera. Primary cells generated from wing biopsies of *R. ferrumequinum*, *M. myotis*, *M. nattereri* and *M. brandtii* (table 1) were subjected to infection by SARS-CoV-2 at a multiplicity of infection (MOI) of 1. Flow cytometry analyses were performed using anti-S antibodies at 24 hours post-infection (hpi). Vero E6 cells, which are African green monkey kidney cells known to be susceptible to SARS-CoV-2 [34] were used as positive controls. Around 40% of Vero E6 cells were positive for the viral S protein (Fig. 1A-B). Neither *R. ferrumequinum* or *Myotis* spp. primary cells were susceptible to SARS-CoV-2 (Fig. 1A-B). We then tested the susceptibility of previously described cell lines generated from *Eptesicus serotinus* [35], *Myotis myotis* [36] and *Tadarida brasiliensis* (table 1) to SARS-CoV-2. *E. serotinus* cells were isolated from brain (FLG) and kidney (FLN) [35](table 1). *M. myotis* cells were established from brain (MmBr), tonsil (MmTo), peritoneal cavity (MmPca), nasal epithelium (MmNep) and nervus olfactorius (MmNol)[36] (table 1). Tb1lu cells are *T. brasiliensis* lung cells. We also generated *Nyctalus noctula* cell lines from lung (NnLu), liver (NnLi) and kidney (NnKi) (table 1). Betacoronaviruses have been sampled in species belonging to these 4 bat genus [8–12]. Human intestinal Caco-TC7 cells and human lung A549 cells, which are both representative of tissues targeted by the virus in infected patients [37], were used as controls. All cells were infected with SARS-CoV-2 at a MOI of 1. Around 23% of Caco-TC7 cells were positive for the viral S protein at 24 hpi (Fig. 1C-D). None of the other selected cells were susceptible to SARS-CoV-2 (Fig. 1C-D).

**Fig 1.**
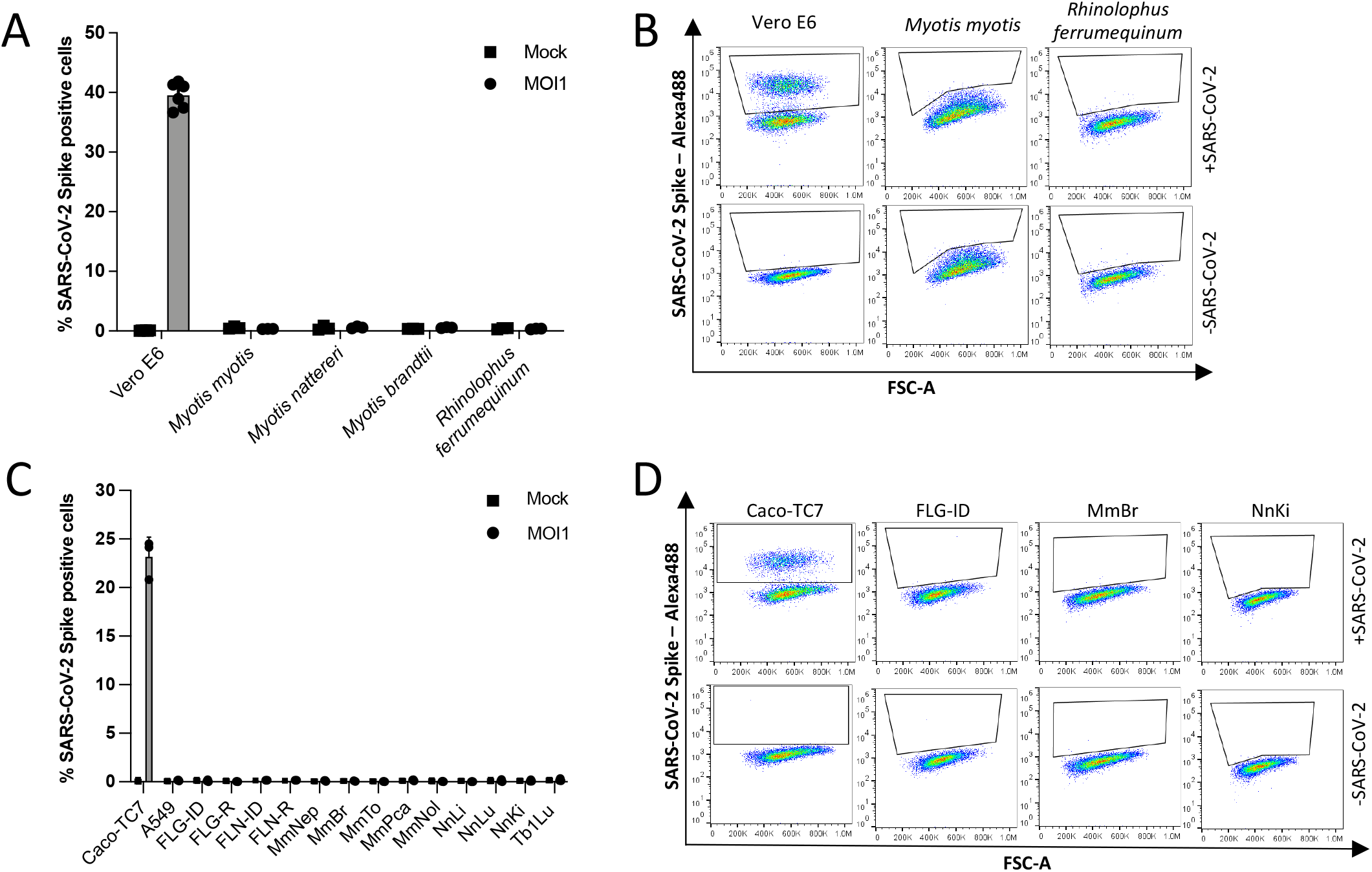
Resistance to SARS-CoV-2 infection in selected bat cell lines. **A**, Primary bat cells derived from wing tissues from four different species, as well as Vero E6 cells, were left uninfected (Mock) or were infected with SARS-CoV-2 at a MOI of 1 for 24 hours and analyzed via flow cytometry for viral spike (S) protein expression. **B**, Representative dot plots of selected cells. Data points represent three technical replicates. **C,** Bat cell lines from four different species, as well as Caco TC7 human intestine and A549 human lung epithelial cells, were left uninfected (Mock) or were infected with SARS-CoV-2 at a MOI of 1 for 24 hours and analyzed via flow cytometry for S expression. **D,** Representative dot plots of selected cells. Data points represent three independent experiments with the exception of A549, FLN-ID and FLN-R cells, where data points represent three technical replicates.

**Table 1.** Overview of bat primary cells and cell lines used in the study.

|  | Bat species | Common name | Family | Organ | Transformation method | Reference |
| --- | --- | --- | --- | --- | --- | --- |
| 16104 | *Rhinolophus ferrumequinum* | greater horseshoe bat | *Rhinolophidae* | Skin (patagium) | None – primary cell | This study |
| 29B | *Myotis myotis* | Common serotine bat | *Vespertilionidae* | Skin (patagium) | None – primary cells | This study |
| 19PL50 | *Myotis nattereri* | Natterer's bat | *Vespertilionidae* | Skin (patagium) | None – primary cells | This study |
| MBra10 | *Myotis brandtii* | Brandt's bat | *Vespertilionidae* | Skin (patagium) | None – primary cells | This study |
| FLG-ID | *Eptesicus serotinus* | Common serotine bat | *Vespertilionidae* | Brain | Immortalized FLG-R cells with SV40 large T antigen | CCLV-RIE 1152 |
| FLG-R | *Eptesicus serotinus* | Common serotine bat | *Vespertilionidae* | Brain | Natural | CCLV-RIE 1093 |
| FLN-ID | *Eptesicus serotinus* | Common serotine bat | *Vespertilionidae* | Kidney | Immortalized FLN-R cells with SV40 large T antigen | CCLV-RIE 1134 |
| FLN-R | *Eptesicus serotinus* | Common serotine bat | *Vespertilionidae* | Kidney | Natural | CCLV-RIE 1091 |
| MmBr | *Myotis myotis* | Greater mouse-eared bat | *Vespertilionidae* | Brain | SV40 large T antigen | He at al., 2014 |
| MmNep | *Myotis myotis* | Greater mouse-eared bat | *Vespertilionidae* | Nasal epithelium | SV40 large T antigen | He at al., 2014 |
| MmNol | *Myotis myotis* | Greater mouse-eared bat | *Vespertilionidae* | Nerve | SV40 large T antigen | He at al., 2014 |
| MmPca | *Myotis myotis* | Greater mouse-eared bat | *Vespertilionidae* | Macrophage | SV40 large T antigen | He at al., 2014 |
| MmTo | *Myotis myotis* | Greater mouse-eared bat | *Vespertilionidae* | Tonsil | SV40 large T antigen | He at al., 2014 |
| NnKi | *Nyctalus noctula* | Common noctule | *Vespertilionidae* | Kidney | SV40 large T antigen | This study |
| NnLi | *Nyctalus noctula* | Common noctule | *Vespertilionidae* | Liver | SV40 large T antigen | This study |
| NnLu | *Nyctalus noctula* | Common noctule | *Vespertilionidae* | Lung | SV40 large T antigen | This study |
| Tb1Lu | *Tadarida brasiliensis* | Mexican/Brazilian free-tailed bat | *Molossidae* | Lung | Natural? | CCLV-RIE 0072 |

SARS-CoV-2 variants exhibiting diverse mutations in the S protein have emerged at the end of 2020, leading to increased transmissibility or/and immune escape in humans [38,39]. The so-called ‘B.1.351/20H/501Y.V2’ and ‘P1/20J/501Y.V3’ variants, which first appeared in South-Africa and Brazil, respectively, have acquired the ability to efficiently replicate in mice airways [40]. We tested the susceptibility of the selected bat cells and Caco-TC7 cells to these two variants. Flow cytometry analysis performed at 24 hpi revealed that none of the bat cell lines were positive for S proteins (Fig. S1A-B). By contrast, the two variants replicated in Caco-TC7 cells (Fig. S1A). Thus, neither the initial virus nor recently emerged variants are able to replicate in the 13 selected bat cell lines.

The lack of production of viral protein in the primary bat cells and bat cell lines, as well as in A549 cells, could be explained by the absence of one or several key pro-viral factor(s) and/or the presence of potent antiviral factor(s).

### Expression of endogenous ACE2 and ectopically-expressed hACE2 in bat cell lines

To determine whether the absence or low expression of ACE2 was the main limiting factor for SARS-CoV-2 replication in the selected bat cell lines, we first evaluated the level of ACE2 expression by RT-qPCR analysis. Levels of ACE2 were above the detection limit in FLG-R, MmTo, MmPca, MmNol cells and in the three Nn cells (Fig. 2A). These cells do not however support viral replication (Fig. 1C-D). Thus, S proteins may have a low affinity for ACE2 expressed in these cells. They may also be deficient in expression of both TMPRSS2 and CTSL. To test this hypothesis, viral input was treated with the serine-protease trypsin to activate the S protein and allow viral fusion in a TMPRSS2- and CTSL-independent manner [19], at the surface of NnKi cells, which express the highest level of ACE2 of all bat cells (Fig. 2A). Trypsin-treated virions did not replicate better than non-treated virions in Caco-T7 (Fig. S1C), suggesting that pre-activation of S proteins does not affect viral fusion in these cells. NnKi cells were resistant to infection with trypsin-treated virions (Fig. S1C), suggesting that S cleavage is not the factor limiting viral infection in these cells.

**Fig. 2.**
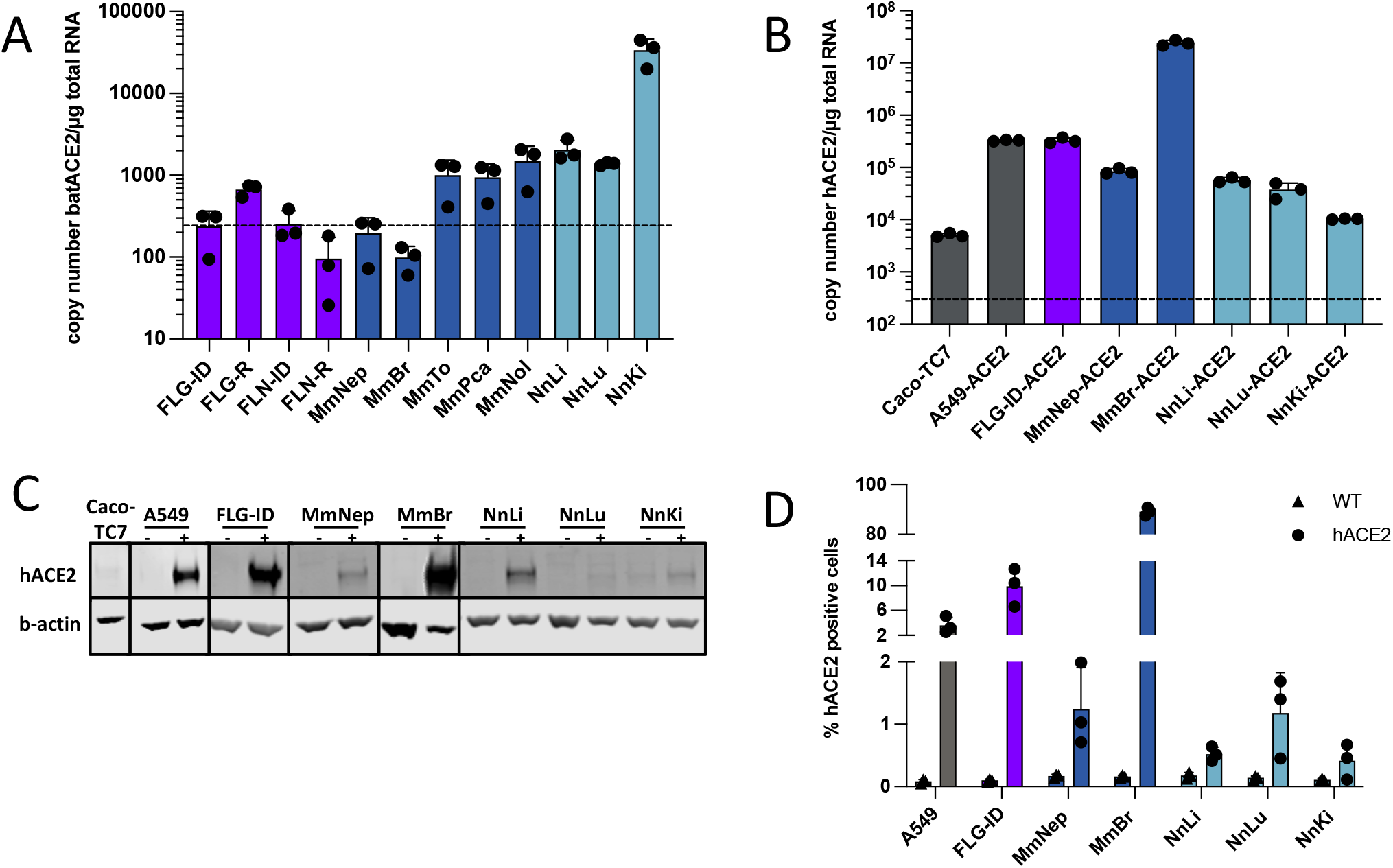
Expression of endogenous ACE2 or ectopically-expressed hACE2 in bat cell lines. **a,** Quantification of copy numbers per μg of total cellular RNA of endogenously expressed ACE2 in indicated bat cell lines via qPCR analysis. **B, C, D,** Indicated bat and human cell lines were stably transduced with a lentivirus vector expressing the hACE2 gene and selected with blasticidin treatment. Human Caco-TC7 intestine cell line served as non-transduced control. **B,** Amount of ectopically-expressed hACE2 gene in each cell line was measured by qPCR analysis and indicated as gene copy number per μg of total cellular RNA. **C,** Whole-cell lysates were analyzed by Western blotting with antibodies against the indicated proteins. Western blots are representative of two independent experiments. **D**, Ectopic hACE2 expression levels of transduced cell lines analyzed via flow cytometry with anti-hACE antibody staining. (a,b,d) Data points represent three independent experiments. (A, B) dotted line indicated limit of detection in qPCR assays.

We then stably expressed hACE2 in bat and A549 cells using lentiviral transduction. Six of the 13 bat cell lines, representing three species (*Myotis myotis*, *Nyctalus noctula* and *Eptesicus serotinus*) tolerated the lentiviral transduction and antibiotic selection. We used RT-qPCR, Western blot and flow cytometry to analyze hACE2 expression in these cell lines. The transduced cells displayed different hACE2 expression profile (Fig. 2B-D). RT-qPCR analysis revealed that hACE2 mRNA abundances were higher in all transduced cells than in Caco-TC7 cells (Fig. 2B), which support SARS-CoV-2 replication (Fig. 1C-D and Fig. S1A). This suggests that transduced cells express hACE2 at a level high enough to permit viral entry. In line with the RT-qPCR analysis, Western blot analysis showed that MmBr-ACE2 cells expressed the highest level of hACE2 among all transduced cell lines (Fig. 2C). ACE2 was barely detectable in Caco-TC7 cells (Fig. 2B). A faint band was also detected in non-transduced NnKi cells, likely representing endogenous bACE2. This suggests that *N. noctula* ACE2 is recognized by the antibody raised against hACE2 in this assay and that Nnki cells expressed higher levels of ACE2 than lung and liver cells from the same bat. These data are in line with the RT-qPCR analysis of endogenous ACE2 expression (Fig. 2A). Flow cytometry analysis revealed that around 80% of MmBr-ACE2 cells and 15% of FLG-ID-ACE2 brain cells were positive for hACE2 (Fig. 2D). On average, 1-2% of A549-ACE2 and MmNep-ACE2 cells were positive for hACE2 and even less Nn cells were expressing hACE2 (Fig. 2D). These low percentages were surprising in light of the RT-qPCR and Western blot data (Fig. 2B and 2C). However, cells counted as negative for hACE2 signal may express levels that are under the detection limit of the assay. Alternatively, anti-ACE2 antibodies may recognize only a subpopulation of the protein by cytometry, such as, for instance, glycosylated and/or truncated forms [41,42]. Of note, endogenous bACE2 expressed in NnKi cells was not detectable in this assay (Fig. 2D).

Despite a potential underestimation of the percentage of hACE2 positive cells by flow cytometry, the three assays revealed that MmBr-ACE2 and FLG-ID-ACE2 cells, both generated from brain tissues, are expressing higher levels of hACE2 than the other transduced bat cell lines. Expression of hACE2 and antibiotic resistance are under the control of 2 different promoters in the bicistronic lentiviral vector we used. Variable strength of the two promoters in the different cell lines could generate cells that survived the antibiotic treatment but express no or very little hACE2. Nevertheless, despite expressing differential levels of hACE2, the hACE2-transduced cells provide models to investigate the interaction between viruses belonging to the SARS-CoV-2 linage and bat cells.

### Expression of hACE2 allows efficient replication of SARS-CoV-2 in *Myotis myotis* and *Eptesicus serotinus* brain cells

The six transduced bat cell lines and A549-ACE2 cells were infected with SARS-CoV-2 for 24 hours at a of MOI of 1. Cytopathic effects (CPEs) were observed in MmBr-ACE2 cells. To illustrate this, we performed time-lapse microscopy of MmBr-ACE2 cells, infected or not, in the presence of propidium iodide (PI) for 48 hours. Cells were rapidly forming syncytia (around 12 hours). Cell death was observed as early as 34 hours, as assessed by the PI uptake through permeable cellular membranes (Fig. 3A-B and movies 1 and 2). Syncytia represent cell-to-cell fusing events mediated by the interaction between cell-surface expressed S proteins and ACE2 [43]. Neither CPE nor syncytia formation were observed in the other cells, as illustrated by the video of infected FLG-ID-ACE2 cells (Fig. 3B-C and movies 3 and 4).

**Fig. 3.**
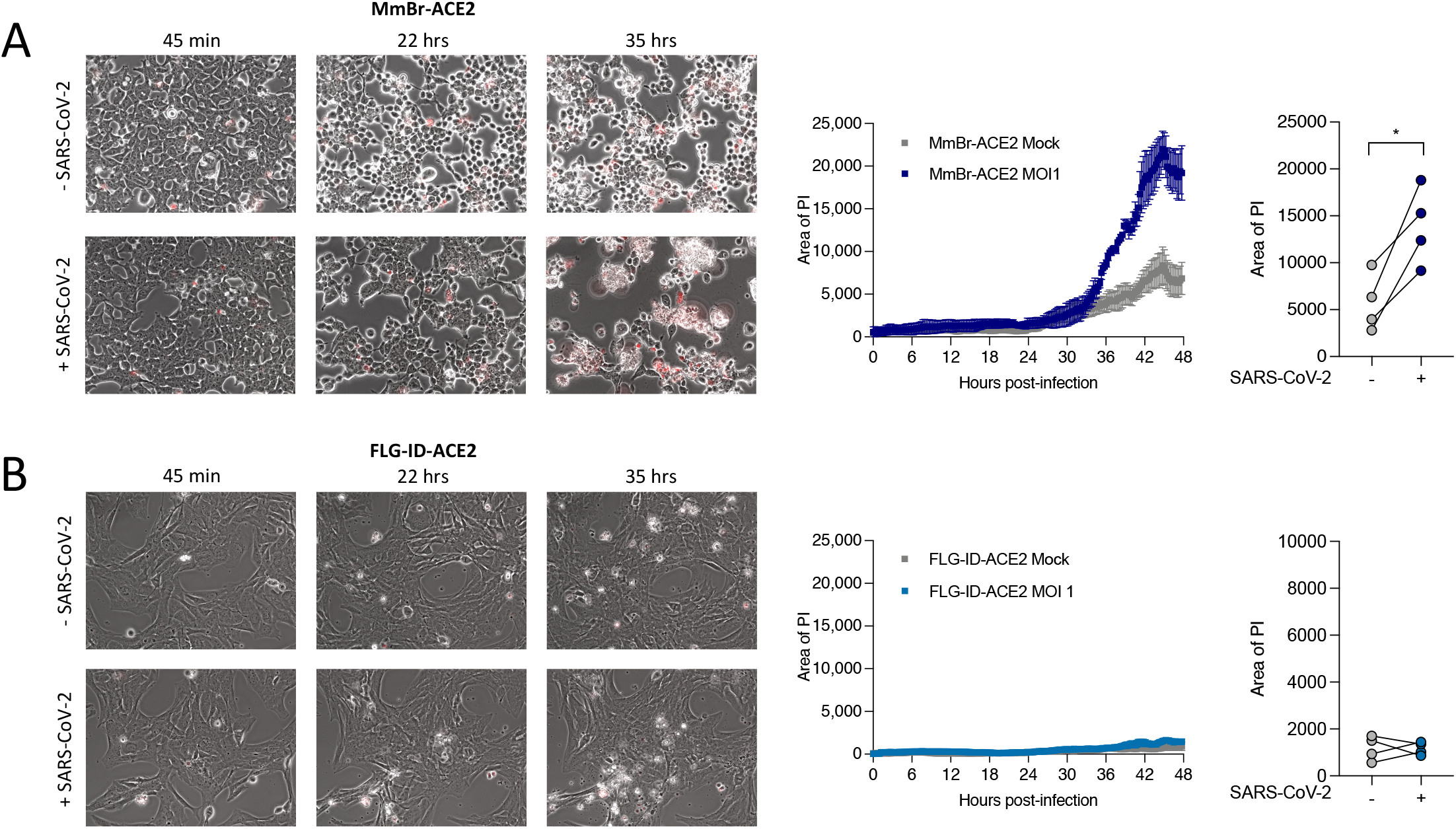
Time-lapse microscopy of *Myotis myotis* and *Eptesicus serotinus* brain cells during SARS-CoV-2 infection. MmBr-ACE2 (**A**) and FLG-ID-ACE2 (**B**) cells were left uninfected (Mock) or were infected with SARS-CoV-2 at a MOI of 1 in media containing propidium iodide (PI) as cell death marker. Images were taken every 10 minutes. Quantification of cell death (area of PI) displayed on the right of corresponding video cutouts. Results are mean ± SD from three fields per condition.

To avoid cell death, MmBr-ACE2 cells were infected with 25 times less viruses (MOI of 0.04) than the other cells (Fig. 4). Assessment of viral replication by RT-qPCR revealed that viral RNA yields increased between 6 and 24 hpi in A549-ACE2 cells, and subsequently reached a plateau (Fig. 4A). Viral RNA yields also increased between 6 and 24 hpi in FLG-ID-ACE2 cells but then dropped back to their 6h-levels (Fig. 4A), suggesting that these cells efficiently controlled viral replication. Viral RNA abundance slightly increased between 6 and 24 hpi in MmNep-ACE2 cells (Fig. 4A), suggesting a low level of viral RNA production, before decreasing at 48 hpi. The profile of viral RNA yield was similar in MmBr-ACE2 cells and in A549-ACE2 cells (Fig. 4A), indicating a robust viral replication; especially, considering that the cells were infected with 25 times less viruses than the others (Fig. 4A). No increase in viral yield was observed in the 3 Nn cell lines between 6 hpi and later time points (Fig. 4A), suggesting an absence of viral replication.

**Fig. 4.**
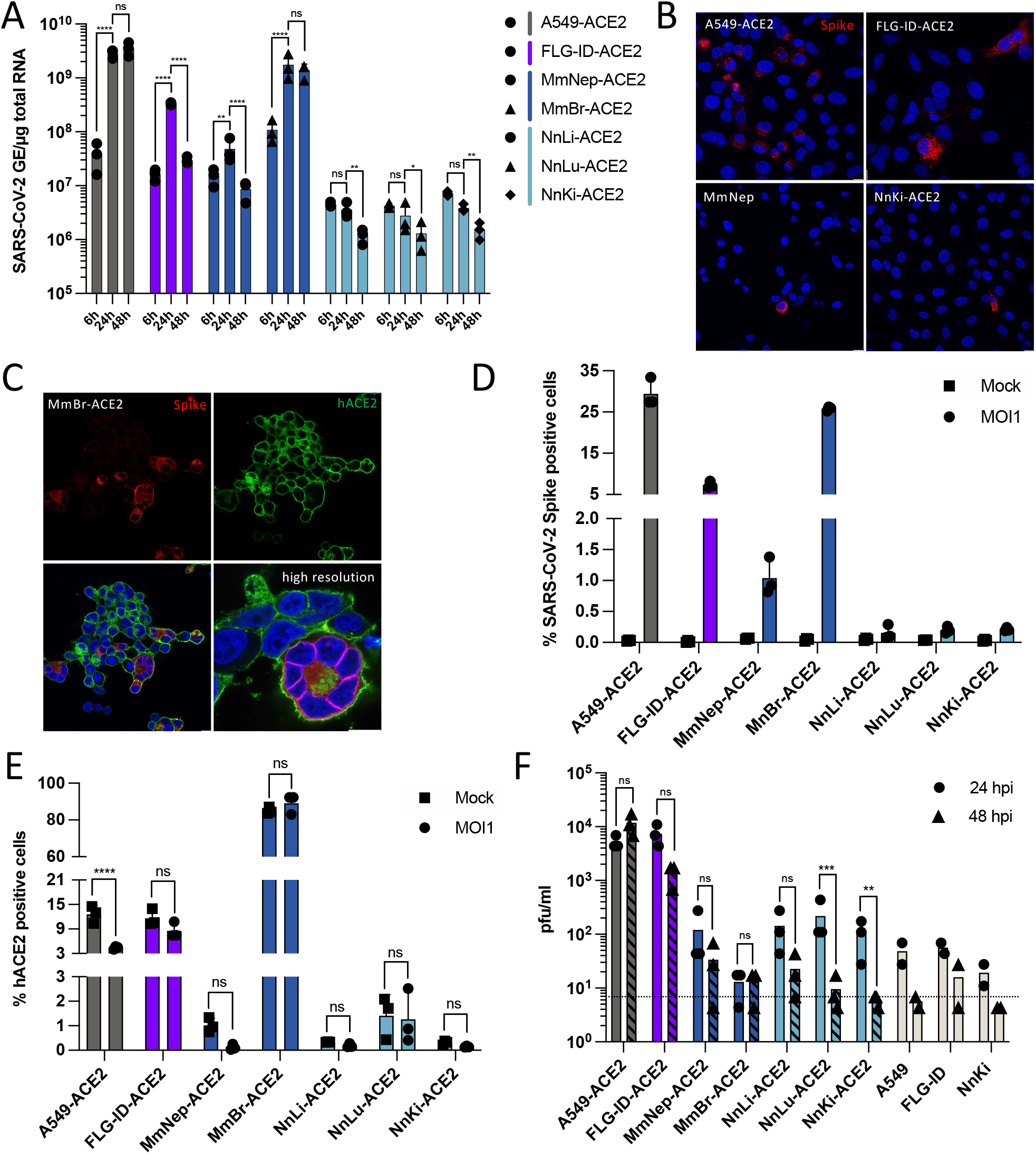
Expression of hACE2 allows efficient replication of SARS-CoV-2 in *Myotis myotis* and *Eptesicus serotinus* cells. Transduced bat cell lines were left uninfected (Mock) or were infected with SARS-CoV-2 at a MOI of 1, with the exception of MmBr cells that were infected at a MOI of 0.04. **A,** The relative amounts of cell-associated viral RNA were determined by qPCR analysis and are expressed as genome equivalents (GE) per μg of total cellular RNA at different time post-infection. All results are expressed as fold-increases relative to uninfected cells. **B, C,** Infected cells were stained at 24 hpi with anti-SARS-CoV-2 S protein (red) and/or anti-hACE2 antibodies (green). Nuclei were stained with Nucblue (blue). Scale bar, 10 μm. **D, E**, The percentages of the indicated cells that contained SARS-CoV-2 S proteins (d) or hACE2 (e) were determined by flow cytometric analysis at 24 hpi. **F,** The presence of extracellular infectious viruses in the culture medium of the indicated cells was determined by TCID_50_ assays with Vero E6 cells at 24 and 48 hpi. Dashed lines indicate the limit of detection. (a, d, e, f) Data points represent three independent experiments. Statistical test: (a) Dunnett’s multiple comparison test on a two-way ANOVA analysis (n.s: not significant; * p-value < 0.05, ** p-value < 0.01, *** p-value < 0.001, **** p-value < 0.0001); (e, f) Šídák’s multiple comparisons test on a two-way ANOVA analysis (n.s: not significant, * p-value < 0.05, ** p-value < 0.01, *** p-value < 0.001, **** p-value < 0.0001).

Cell lines that seemed to support viral replication (Fig. 4A), as well as one Nn cell line, were analyzed for the expression of viral proteins through immunofluorescence imaging using antibodies specific for S and for hACE2 at 24 hpi. Cells positive for S were observed in all cell lines (Fig 4B, C). However, the proportion of positive cells varied considerably between them (Fig 4B), confirming disparities in viral susceptibilities between cells of different species and/or tissues (Fig. 4A). For instance, almost no cells were expressing the S protein in NnKi cells (Fig. 4B). An hACE2 signal was only detected in MmBr-ACE2 cells (Fig. 4C), which are the cells that express the most hACE2 among the transduced cell lines (Fig. 2B-D). Thus, as previously observed in flow cytometry assays (Fig. 2D), the selected anti-hACE2 antibody appeared to allow detection in immunofluorescence analysis only when the protein is expressed at high levels. The confocal images also confirmed the presence of syncytia in MmBr-ACE2 infected cells (Fig. 4C). To quantify the disparities in viral protein production between cells, flow cytometry analyses were performed. On average 25% of A549-ACE2 and MmBr-ACE2 cells were positive for S protein when infected for 24 hours at a MOI of 1 or 0.04, respectively (Fig. 4D). Around 5% of FLG-ID-ACE2 and 1% of MmNep-ACE2 cells were expressing the S protein (Fig. 4D). Less than 0.2% of Nn cells were positive for the S protein (Fig. 4D). These flow cytometry data agree with both viral RNA yields (Fig. 4A) and e immunofluorescence analysis (Fig. 4B-C). The same samples were stained with anti-hACE2 antibodies. Only around 3% of infected A549-ACE2 cells appeared hACE2-positive (Fig. 4E) while around 25% of them were S-positive (Fig. 4D). Knowing that these cells are not permissive to viral replication in the absence of hACE2 over-expression (Fig. 1A), these results further suggest that the anti-hACE2 antibodies recognized only a subpopulation of ACE2. Similarly, only a fifth of FLG-ID-ACE2 cells were double-positive (Fig. S2A). This under-estimation of hACE2 positive cells could also be explained by the presence of S-induced syncytia (Fig. 3a and 4c), which are indeed detectable using the forward (FSC) and sideward scatter (SSC) parameters of the cytometer (Fig. S2B), and likely affects the cell count. ACE2 expression has also been reported to be downregulated in infected human intestinal organoids [44]. This is also the case for infected A549-ACE2 cells (Fig. 4E).

Virus titration on Vero E6 cells showed that A549-ACE2 cells released around 6.10^3^ PFU/ml and 10^4^ PFU/ml at 24 and 48 hpi, respectively (Fig. 4F). Despite producing less viral RNA than A549-ACE2 cells, FLG-ID-ACE2 cells yielded similar amounts of infectious particles at 24hpi (Fig. 4F). Albeit not significant, less infectious particles were produced from FLG-ID-ACE2 cells at 48 hpi than at 24 hpi (Fig. 4F), which is in accordance with a decrease of viral RNA production between 24 hpi and 48 hpi (Fig. 4A). These data further suggest that viral replication is controlled in *E. serotinus* brain cells. As expected from viral RNA and viral protein quantification (Fig. 4A and 4B), MmNep-ACE2 cells produced only small amounts of infectious particles, around 100 PFU/ml at 24 hpi and around 60 PFU/ml at 48 hpi (Fig. 4F). MmBr-ACE2 cells released only around 10 PFU/ml, which is 1000 times less than A549-ACE2 cells (Fig. 4F). This was surprising since the two cell lines produced similar quantities of viral RNAs and proteins (Fig. 4A-D). Approximately 100 PFU/ml were collected from the supernatant of the three lines of Nn cells, a similar amount to what was detected in non-permissive cells, such as non-transduced A549, FLG-ID and Nnki cells, which were included in the analysis as negative controls (Fig. 4F). These infectious particles are thus likely input viruses that were carried over from the inoculum of the first round of infection.

Together, these data revealed that expression of hACE2 allowed the virus to complete its replication cycle in *E. serotinus* FLG-ID brain cells, suggesting an ACE2-mediated refractory state to SARS-CoV-2 replication. Expression of hACE2 in *M. myotis* brain cells (MmBr-ACE2) allows the production of viral RNA and proteins, indicating that the ACE2-mediated restriction can also be overcome. However, infectious particles were not released from these cells, suggesting the existence of another cellular restriction at a later stage of the viral replication cycle. In MmNep-ACE2 and Nn cells, expression of hACE2 was not sufficient to allow robust viral replication, suggesting a deficiency in key proviral factor(s) and/or expression of potent antiviral factor(s).

### Infectious particles are produced by MmBr-ACE2 cells but are not released

Since MmBr-ACE2 cells sustained the production of viral RNAs and proteins (Fig. 4A-C-D), we were intrigued by the absence of infectious particles release (Fig. 4F). Despite infecting these cells with a MOI of 0.04 (Fig. 4) to reduce the CPEs observed at a MOI of 1 (Fig. 3), we wondered whether cytokines released by infected cells and/or dying cells may stimulate damage-associated molecular patterns (DAMPs) and thus trigger an antiviral response inhibiting viral replication in Vero E6 cells. Other possibilities include a defect in viral assembly and/or in viral transport through the secretory pathway in MmBr-ACE2 cells. Alternatively, these cells may only produce immature non-infectious viral particles. To investigate these hypotheses, supernatants collected from MmBr-ACE2 cells, and as controls, from A549-ACE2 and MmNep-ACE2 cells, were clarified by ultracentrifugation to get rid of potential cytokines and cell debris and titrated on Vero E6 cells. Flow cytometry analysis using anti-S antibodies were done on the same samples to verify that the cells were infected (Fig. 5A). Clarified and ultracentrifuged supernatants from A549-ACE2 and MmNep-ACE2 cells contained similar amounts of infectious particles, around 10^4^ PFU/ml and 100 PFU/ml, respectively (Fig. 5B). Comparable to observations in previous experiments (Fig. 4F), little infectious particles, around 10 PFU/ml, were recovered in both the clarified and ultracentrifuged supernatant of infected MmBr-ACE2 cells (Fig. 5B). These data suggest that immunostimulatory components, such as cytokines or dying cells, that could be present in infected MmBr-ACE2 cell culture supernatants did not affect the results of the titration assays. To assess the presence of intracellular infectious particles in MmBr-ACE2 cells, the titration assays were performed on crude cell lysates collected at 24 hpi. Around one log more infectious particles were retrieved from lysed A549-ACE2 cells than from their supernatant (Fig. 5B). By contrast, only 10^2^ PFU/ml viral particles were collected in lysed MmNep-ACE2 cells (Fig. 5B). These results agree with the level of viral replication previously detected in A549-ACE2 and MmNep-ACE2 cells (Fig. 4). Around 3 log more viral particles (about 10^4^ PFU/ml) were retrieved from lysed MmBr-ACE2 cells than in the culture supernatant (Fig. 5B), suggesting that viral assembly and maturation takes place in these cells and that the absence of viral release is likely due to a defect in viral transport through the secretory pathway. Thus, MmBr-ACE2 cells are either missing one or several cellular factor(s) that are required for exit of infectious virions from assembly sites to the cell membrane and/or they express one or several antiviral factor(s) that potently block this transport.

**Fig. 5.**
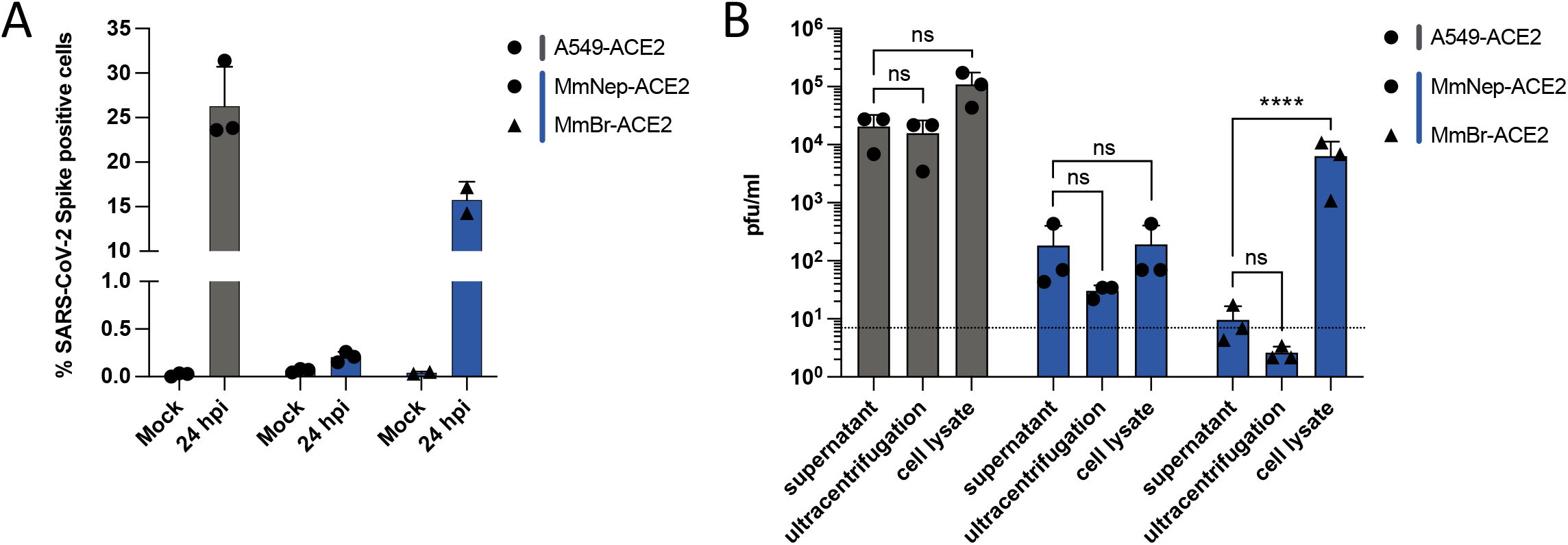
Infectious particles are produced by MmBr-ACE2 cells but not released. A549-ACE2 and MmNep-ACE2 cells were left uninfected (Mock) or were infected at a MOI of 1 for 24 hours. MmBr-ACE2 cells were left uninfected (Mock) or were infected at a MOI of 0.04 for 24 hours. **A,** The percentages of the indicated cells that contained SARS-CoV-2 S proteins were determined by flow cytometric analysis. **B,** The presence of extracellular infectious viruses in the culture medium of the indicated cells was determined by TCID_50_ assays performed on Vero E6 cells. Supernatants were either clarified or purified by ultracentrifugation. Alternatively, cell-associated infectious virions were titrated on Vero E6 cells from whole cell lysates. Data points represent three independent experiments. Statistical test: Dunnett’s multiple comparison test on a two-way ANOVA analysis (n.s: not significant, * p-value < 0.05, ** p-value < 0.01, *** p-value < 0.001, **** p-value < 0.0001).

### An abortive entry route exists in bat and human cells

To investigate further ACE2-mediated restriction, we performed binding and entry assays on cells transduced or not with hACE2 (Fig. 6A). Infected cells were kept on ice for 1 hour, washed three times and then either lysed (‘on ice’) or incubated at 37 degrees for 2 or 6 hours. To remove potential residual bound particles, the warmed cells were treated with trypsin for 30 minutes prior to lysis. We performed the assays with A549, FLG and NnKi cells since they tolerated the three washes on ice without detaching from the plates and, for each cell line, we compared viral RNA abundance in wild-type *versus* hACE2-expressing cells. Viral RNA detected in cells that were kept on ice represent input viruses bound to cellular membranes. In all six cell lines, we indeed detected viral RNA bound to cell membranes (Fig. 6A), suggesting that hACE2 expression is not required for viral attachment. Such ACE2-indepenent binding of the S protein could be mediated by heparan sulfate, as described for several human cell lines [45,46], or by endogenous ACE2 when it is expressed at detectable levels (Fig. 2A). hACE2 expression may however enhance viral binding to the A549 and FLG cell membranes since around 500 more genome copies per μg of total RNA were detected in cold transduced cells than in unmodified ones (Fig. 6A). Abundance of viral RNAs increased between 2 and 6 hours both in A549-ACE2 and FLG-ACE2 cells but not in wild-type cells (Fig. 6A). These results confirm that viral replication occurred hACE-2 expressing A549 and FLG cells (Fig. 4). No increase in viral RNA yield was observed between 2 and 6 hours in NnKi-ACE2 cells (Fig. 6A), confirming the absence of viral replication in these cells (Fig. 4). Viral RNA detected at 2 or 6 hpi in non-transduced cells (Fig. 6A) may represent viruses that remained attached to the cell surface despite the trypsin treatment or viruses that penetrated the cells via an hACE2-independent route. To ensure that the trypsin treatment was effective in cleaving off particles bound to the cell surface, A549 and NnKi cells kept on ice for one hour were treated with trypsin for 30 minutes (Fig. 6B). Around 2 to 3 log less viral RNA was detected in trypsinized A549 and NnKi cells than in non-treated cells (Fig. 6B), suggesting that a large quantity of viruses is indeed detaching from cell membranes upon trypsin treatment. Significantly more viral RNA was detected in A549 and NnKi cells that had been shifted to 37 degrees for 2 h than in iced cells treated with trypsin (Fig. 6B). These viral RNA molecules may represent virions that penetrated the cells. Together, these data suggest that viruses are internalized in cells that do not over-express ACE2 (Fig. 6). This internalization path does not, however, lead to a productive viral cycle (Fig. 1 and 4).

**Fig. 6.**
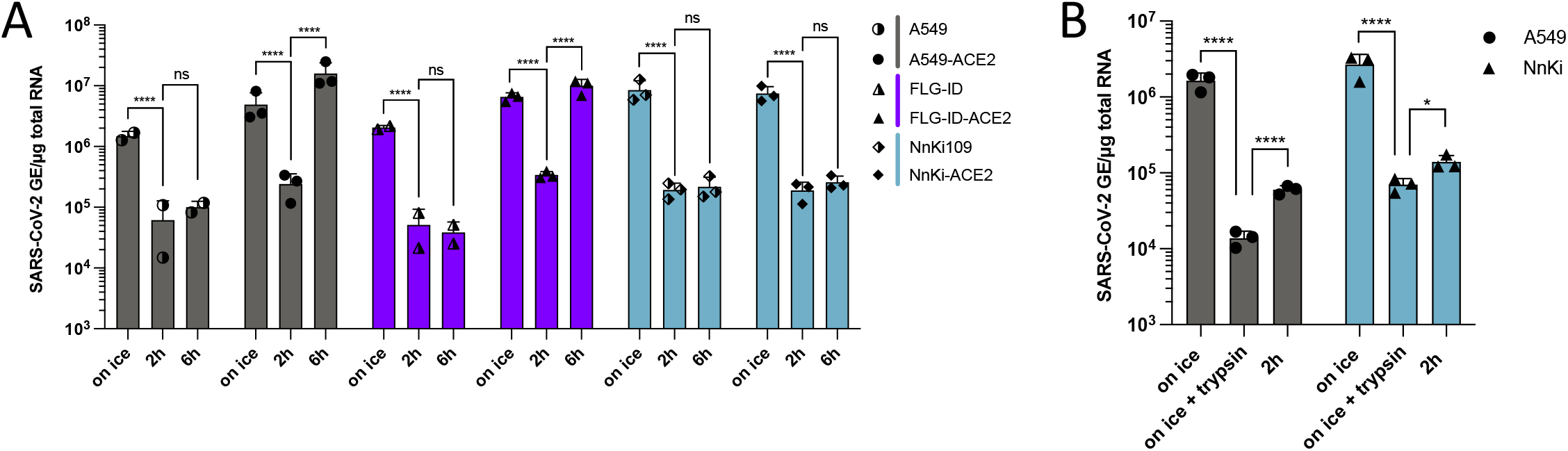
An abortive entry route exists in bat and human cells. **A,** Cells were incubated with SARS-CoV-2 at a MOI of 1 for 1 hour on ice to allow viral attachment. After extensive washing, a portion of the cells was lysed (“on ice”) and the remaining cells were incubated for 2 or 6 hours at 37°C to permit viral internalization. After the incubation period, these cells were lysed after 30 min trypsinization to remove bound viruses from the cell surface (“2h”, “6h”). **B,** A 30 min trypsinization step was added after the initial incubation on ice (“on ice + trypsin”). The “on ice” and “2h” conditions are the same as (a). **A, B** The relative amounts of cell-associated viral RNA were determined by qPCR analysis and are expressed as genome equivalents (GE) per μg of total cellular RNA. Data points represent three independent experiments. Statistical test: Dunnett’s multiple comparison test on a two-way ANOVA analysis (n.s: not significant, * p-value < 0.05, ** p-value < 0.01, *** p-value < 0.001, **** p-value < 0.0001).

### Viral IFN counteraction mechanisms are species-specific

Quantification of intracellular viral RNAs and titration assays revealed that FLG-ACE2 cells, and, to a lesser extent, MmNep-ACE2 cells, controlled viral replication over time (Fig. 4A and D). By contrast, viral RNA yield remained high between 24 and 48 hpi in A549-ACE2 and MmBr-ACE2 cells (Fig. 4A and D). To assess whether the interferon (IFN) response could contribute to viral containment in FLG-ACE2 and MmNep-ACE2 cells, we compared mRNA abundance of two IFN-stimulated genes (ISGs) upon stimulation or infection in the different cell lines. We selected OAS1 and IFIH1, 2 ISGs that are conserved across vertebrate species [47]. Moreover, OAS1 expression is associated with reduced COVID-19 death [48] and IFIH1 codes for Mda5, the protein responsible for sensing SARS-CoV-2 replication intermediates, and thus initiating the IFN response, in human cells[49,50]. We first evaluated the expression of the two selected ISGs upon transfection with polyI:C, a synthetic dsRNA analog. All seven cell lines contained transcripts for these two ISGs and responded well to the stimulation (Fig. 7A-B), demonstrating that they possess intact IFN-induction and -signaling pathways.

**Fig. 7.**
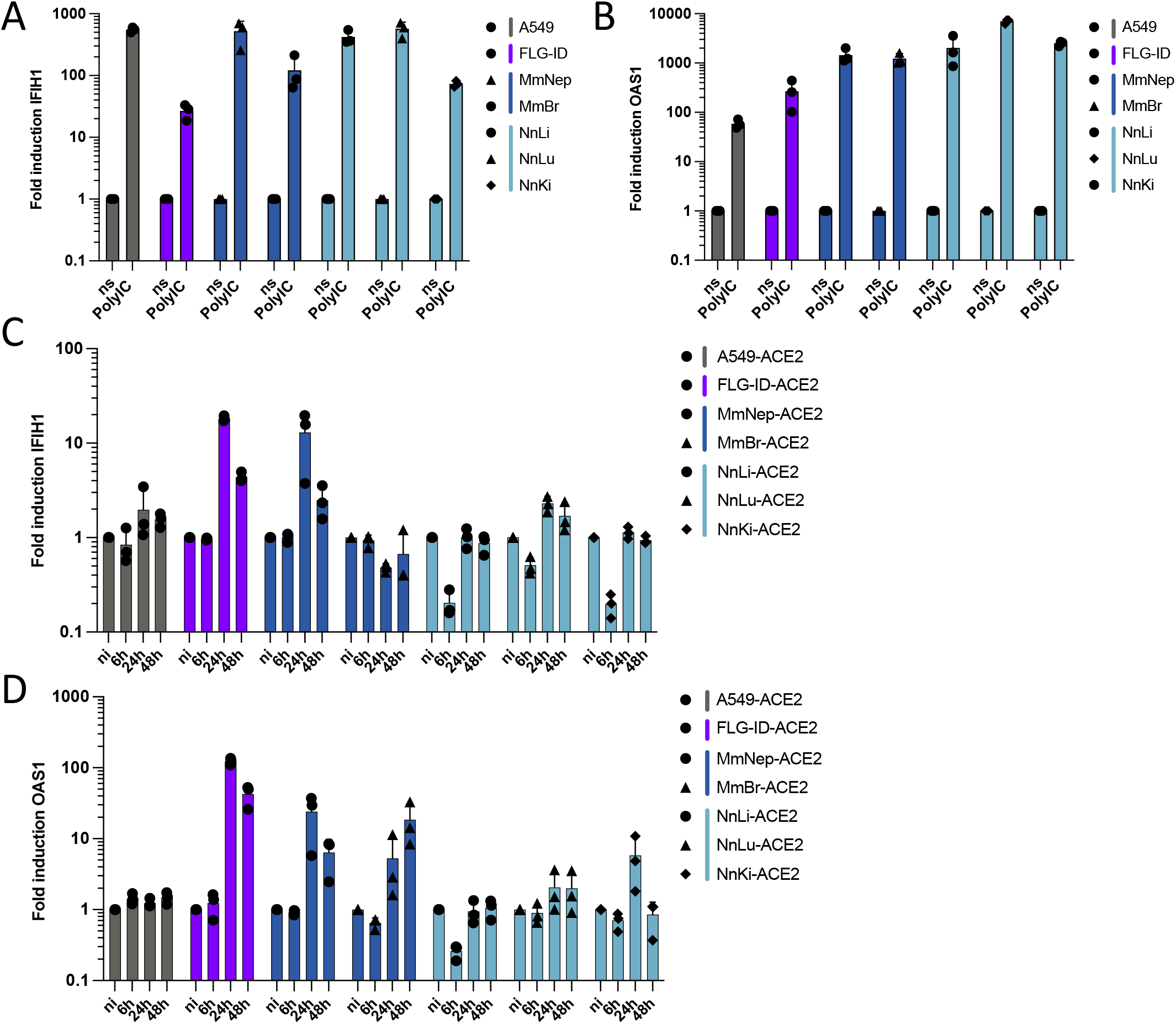
Viral IFN counteraction mechanisms are species-specific. **A, B,** Non-transduced cell lines were transfected with 250 ng low-molecular weight PolyI:C or were treated with PBS for 16 hours. The relative amounts of *IFIH1* mRNA (a) and *OAS1* mRNA (b) were determined by qPCR analysis. Results are expressed as fold-increases relative to unstimulated PBS-treated cells. **C, D**, Whole cell lysates of infected cells (same lysates used for viral quantification in panel 4a) were analyzed via RT-qPCR assays for the relative amounts of *IFIH1* mRNA (c) and *OAS1* mRNA (d). Results are expressed as fold-increases relative to uninfected cells. (**A-D**) Glyceraldehyde 3-phosphate dehydrogenase (GAPDH) of corresponding species was used as house-keeping gene. Data points represent three independent experiments.

We then evaluated the mRNA abundance of these two ISGs in cells infected for 6, 24 and 48 hours (Fig. 7C-D). No increase of OAS1 and IFIH1 expression was observed in A549-ACE2 cells (Fig. 7C-D). This agrees with a previous report showing that infection of A549-ACE2 by SARS-CoV-2 is characterized by an absence of IFN response [51]. By contrast, the abundance of OAS1 and IFIH1 transcripts increased between 6 and 24 hpi in FLG-ACE2 and MmNep-ACE2 cells (Fig. 7C-D) and remained elevated at 48 hpi in both cell lines (Fig. 7C-D). In MmBr-ACE2 cells, the infection triggered the induction of OAS1 expression but not of IFIH1 (Fig. 7C-D), suggesting that the virus is able to suppress IFN-mediated Mda5 upregulation in this cell type. No or little stimulation of OAS1 and IFIH1 expression was observed in Nn cells upon viral infection (Fig. 7C-D). This was expected since the virus replicates at very low levels in these cells (Fig. 4). Surprisingly, the mRNA abundances of both OAS1 and IFIH1 were lower in NnLi-ACE2 cells exposed to the virus for 6 h than in control cells (Fig. 7C-D). Such downregulation was also observed for IFIH1 in NnKi-ACE2 cells. Since only around 0.1% of Nn cells are producing S proteins (Fig. 4D), this decrease in OAS1 and IFIH1 expression is replication-independent. It could be mediated by innate immune sensors present at the cell surface and/or in endosomal compartments, where the virus may be retained (Fig. 6), such as toll-like receptors (TLRs) [52].

Together, our data confirmed that SARS-CoV-2 efficiently counteracts ISG induction in A549-ACE2 cells [51] and revealed that it is not the case in FLG-ACE2 and MnNep-ACE2 cells. The control of viral replication observed in these two cell lines (Fig. 4) could thus be due to the expression of a set of ISGs with potent antiviral functions. Interestingly, IFN-mediated barriers are not only species-specific but also organ-specific since the virus dampens Mda5 expression in MmBr-ACE2 cells but not in MmNep-ACE2 cells.

## Discussion

The development of novel bat cellular models is essential to understand the molecular mechanisms underlying the ability of bats to serve as reservoirs for numerous viruses, including alpha- and beta-coronaviruses. We first produced *R. ferrumequinum*, *M. myotis*, *M. nattereri* and *M. brandtii* primary cells to evaluate their susceptibility to infection with the initial SARS-CoV-2. None of them supported viral replication, not even *R. ferrumequinum*, which belongs to the same genus as the host (*R. affinis*) of RaTG13, a potential ancestor of SARS-CoV-2 [4]. These primary cells, which were generated from patagium biopsies of living bats, exhibited a dermal-fibroblast phenotype. A single-cell transcriptomic analysis showed that *R. sinicus* skin cells express moderate levels of ACE2 and very little TMPRSS2 [53]. The virus may thus be able to enter the skin primary cells that we generated but the fusion step could be the factor limiting infection. Further experiments will be required to characterize at which step of its replicative cycle the virus is stopped in these primary cells.

We established the first three *Nyctalus noctula* cell lines using liver, kidney and lung tissues from a single bat. These organs are site of viral replication in infected patients [37] and may thus be physiologically relevant for bat infection as well. Similar to the cells originated from *M. myotis* and *E. serotinus* bat species, all three Nn cell lines responded well to stimulation with synthetic dsRNA, indicating that they are valuable tools to study the bat innate immune response. In addition to coronaviruses, *N. noctula* carries other viruses with zoonotic potential such as paramyxoviruses and hantaviruses [54,55]. The Nn cells that we have developed represent thus novel opportunities to study bat-borne viruses. We found that Nn kidney cells expressed higher levels of ACE2 than Nn cells derived from lung or liver. Likewise, ACE2 is expressed at high levels in *R. sinicus* kidney, as revealed by comparative single-cell transcriptomic [53] and *in silico* [56] analysis of ACE2 expression pattern in various tissues. ACE2 is also highly expressed in human kidney [57]. Thus, kidney cells appear relevant to study betacoronavirus replication. Finally, we have generated six bat cell lines expressing hACE2. The varying susceptibilities of the six transduced cells to SARS-CoV-2 infection offer opportunities to decipher species-specific antiviral mechanisms that have evolved in bats. A obvious need to develop additional bat cell lines still remains [30]. Particularly valuable cells would be cells derived from bat intestine, a tissue that expresses high level of proteins known to mediate or facilitate cellular entry of bat-borne betacoronaviruses, such as ACE2 and TMPRSS2, at least in *R. sinicus* [53], and that is relevant for SARS-CoV-2 infection, as demonstrated by the detection of viral genomes in duodenum tissue of experimentally infected *Rousettus aegypticus* [29].

*Myotis myotis*, *Tadarida brasiliensis*, *Eptesicus serotinus*, and *Nyctalus noctula* cells were resistant to infection with both the initial virus and two recently emerged variants. ACE2 from *M. myotis* and *T. brasiliensis*, as well as from a species of the *Eptesicus* genus, permitted S-mediated entry of pseudotyped VSV when ectopically expressed in human cells refractory to SARS-CoV2 infection [22]. This means that when expressed at high levels, ACE2 from these three species interacts with the viral S protein. As in human A549 cells, ACE2 may be expressed at a level which is too low to allow viral entry in our bat cell line models. Potential ability of *N. noctula* ACE2 to bind S protein has not been reported and the genome of this bat genus is yet to be sequenced. Hence, low affinity between S protein and ACE2 and/or low level of ACE2 expression may hamper viral replication in these cells. Our results highlight the importance of performing experiments in the context of genuine infection of bat cells to predict their susceptibility to infection.

Trypsin-resistant viruses were detected in non-transduced A549, FLG-ID and NnKi cells at early time post-infection. They likely represent input viruses that penetrated the cells despite the absence of hACE2 expression. These viruses could have entered cells using bACE2 or via an ACE2-independent manner. Such ACE2-independent entry has been described previously in Vero E6 cells [19]. By contrast to what is observed in Vero E6 cells, these two potential paths are abortive in A549, FLG-ID and NnKi cells since it doesn’t allow viral replication. The virus may be routed to a subset of endosomes that lack appropriate proteases. Expression of hACE2 allowed efficient viral RNA and protein production in A549, FLG-ID and Mm cells, suggesting that these cells express proteases that efficiently cleave S proteins. It also shows that ACE2 alone was responsible for the lack of viral replication in non-transduced A549, FLG-ID and Mm cells. This ACE2-mediated entry block is rather due to a low or absent ACE2 expression than to an incompatibility between ACE2 and S protein since ectopic expression of *Myotis* spp. and *Eptesicus* spp. ACE2 facilitate S-mediated entry of pseudo-viruses [22,23]. By contrast, expressing hACE2 in Nn cells, at a higher level than in permissive Caco-TC7 cells, seems sufficient to permit internalization but not replication. Nn cells may not express proteases able to cleave S protein nd virions are thus probably retained in endosomes.

Infectious particles were produced in Mm cells but were not released into the extracellular milieu. Instead of using the canonical secretory pathway exploited by many enveloped viruses to exit cells, betacoronaviruses hijack lysosomes for their transport from assembly sites to the plasma membrane [58]. Mm cells maybe deficient in one or several components of this lysosomal pathway. Since infection induced S-mediated syncytia formation in Mm cells, viruses might spread from cell-to-cell via syncytia, as do other syncytia-forming viruses such as respiratory syncytial virus, parainfluenza viruses and measles[59]. Syncytia-mediated intercellular spreading allows viruses to escape virus-neutralizing antibodies. Such mode of transport has been previously proposed in human cells infected with the Middle East Respiratory Syndrome coronavirus (MERS-CoV) [60], another betacoronavirus. Analysis of *post-mortem* samples of patients that succumb of COVID-19 revealed the presence of syncytial pneumocytes positives for viral RNAs [61]. However, the pathogenetic significance of syncytia remains to be investigated.

SARS-CoV-2 has evolved numerous synergetic mechanisms to evade the IFN response in human cells [62], resulting in an absence of IFN expression in some cells, including A549 cells [51,63]. The virus is unable to counteract ISG induction in *Eptesicus serotinus* kidney cells and in *Myotis myotis* nasal epithelial cells. This is especially intriguing in *E. serotinus* cells since the virus replicates to high levels in these cells and thus produce proteins with described IFN antagonist activities. Similarly, MERS-CoV suppresses the antiviral IFN response in human cells but not in *E. fuscus* cells [64]. One can envisage that escape of IFN-mediated restriction by betacoronaviruses is species-specific. For instance, SARS-CoV-2 Nsp14 targets human IFNAR1 for lysosomal degradation [62], but may be unable to degrade bat IFNAR1. This inability to evade IFN response in *Eptesicus serotinus* kidney cells and in *Myotis myotis* nasal epithelial cells may contribute to the cellular control of infection in FLG-ID and MmNep cells, as in experimentally infected *Eptesicus fuscus* [65]. Other mechanisms could be at play. For instance, the basal level of IFN may be high in these two cell lines, as reported in several other bat species[66–68]. Expression of a mutated form of IRF3, which is a key transcription factor involved in the induction of the IFN signaling cascade, contributes to an enhanced IFN response in bat species, including in *E. fuscus*, as compare to human [69]. Investigation of IRF7, another transcription factor that mediates IFN expression, in *Pteropus alecto* cells revealed a more widespread tissue distribution in bats than in humans [70,71]. Bats may thus launch IFN-dependent measures against viruses in a faster and broader manner than in humans [72]. Another possibility to explain the high level of ISG expression in infected *E. serotinus* kidney cells is that they express specific set of potent antiviral ISGs. Expression of atypical ISGs has been reported for different bat species, including RNA-degrading ribonuclease L (RNaseL) in *P. alecto* cells and RNA-binding Microrchidia 3 (MORC3) in *Pteropus vampyrus* and *Eidolon helvum* cells [66,73]. Pursuing the characterization of bat innate immunity in relevant *in vitro* models is essential to understand the mechanisms by which they control the replication of numerous unrelated viruses.

## Materials and Methods

### Bat primary cells

*M. myotis* samples were collected in July 2020 from two bat colonies in Inca and Llucmajor on Mallorca (Balearic Islands, Spain) (agreement CEP 31/2020 delivered by the Ministry of the Environment and Territory, government of the Balearic Islands). *R. ferrumequinum* biopsies were collected in France in 2020. Authorization for bat capture was delivered by the French Ministry of Ecology, Environment and Sustainable development (approval C692660703 from the Departmental Direction of Population Protection (DDPP), Rhone, France). All methods were approved by the ‘Muséum National d’Histoire Naturelle (MNHN)’ and the ‘Société Française pour l’Étude et la Protection des Mammifères (SFEPM)’. Patagium biopsies were shipped in freezing medium Cryo-SFM (PromoCell), on dry ice or at 4°C with ice packs. Primary cells were obtained as previously described [74,75]. Briefly, skin biopsies were washed twice with sterile PBS, excised in small pieces and enzymatically digested, either with 500 μL of collagenase D (1 mg/mL) (Roche) and overnight incubation at 37°C without agitation, or with 100-200 μL of TrypLE Express Enzyme (Gibco) and incubation 10 min at 37°C under gentle agitation. Dissociated cells and remaining pieces of tissue were placed in a single well of a 6-well plate containing 2 mL of Dulbecco’s Modified Eagle Medium (DMEM, Gibco) containing 20% heat-inactivated fetal bovine serum (FBS) (Eurobio), 1% penicillin/streptomycin (P/S) (Gibco), and 50 μg/ mL gentamycin (Gibco), and incubated at 37°C under 5% CO_2_. Cell cultures were regularly checked to determine the need for media refreshment or splitting. After 5-10 passages, cells were grown in DMEM supplemented with 10% FBS.

### Cell lines

FLG-ID, FLG-R, FLN-ID, FLN-R and Tb1Lu cell lines (table 1) were maintained in equal volumes of Ham’s F12 and Iscove’s modified Dulbecco’s medium (IMDM, Gibco), supplemented with 10% FBS and 1% P/S (Gibco) in non-vented flasks. Mm cells, which were obtained from a single common serotine bat (*Eptesicus serotinus*), were previously described [36]. Nn kidney-, liver- and lung-derived cell cultures were obtained from a common noctule bat (*Nyctalus noctula*) euthanized because of poor chance of survival associated with traumatic injuries sustained while a dead tree sheltering bat hibernaculum was cut. The decision to euthanize the specimen was made by a veterinarian following inspection of a group of noctule bats presented for examination and therapy in the rescue center at the University of Veterinary and Pharmaceutical Sciences Brno, Czech Republic, in November 2015 [76]. The bat was anesthetized with isofluranum (Piramal Enterprises Ltd.) and euthanized by quick decapitation. The cadaver was immersed into 96% ethanol for a few seconds and then subjected to necropsy under aseptic conditions to collect organs which were loosened mechanically with scalpel blades, minced into small pieces, suspended in DMEM (Biosera) containing 1 mg/ml collagenase (Thermo Fisher Scientific) and 1 mg/ml trypsin (Sigma-Aldrich), and incubated at 37 °C on a shaking thermoblock for 45 min. The cells were then separated through a 100 μm nylon filter and washed twice in a medium supplemented with 10% FBS to stop enzymatic digestion. The cells yielded in this way were cultured in DMEM supplemented with 10% FBS and 1% P/S (Sigma). Primary cells were immortalized by transfection of pRSVAg1 plasmid expressing Simian Vacuolating Virus 40 large T antigen (SV40T) with lipofectamine 2000 (Invitrogen) according to the manufacturer’s protocol, expanded and cryopreserved. Mm and Nn cell lines (table 1), as well as African green monkey Vero E6 cells (ATCC CRL-1586), human lung epithelial A549 cells (kind gift from Frédéric Tangy, Institut Pasteur, Paris) and human colorectal adenocarcinoma Caco TC7 cells (ATCC HTB-37), were maintained in DMEM (Gibco), supplemented with 10% FBS and 1% P/S in vented flasks. All cells were maintained at 37°C in a humidified atmosphere with 5% CO_2_. Bat and A549 cells were modified to stably express hACE2 using the pLenti6-hACE2 lentiviral transduction as described previously [43]. Briefly, 2×10^5^ cells were resuspended in 150 μl of culture medium containing 15 μl of ultracentrifuged lentiviral vectors supplemented with 2mM HEPES (Gibco) and 4 μg/ml polybrene (Sigma). Cells were agitated for 30 sec every 5 min for 2.5 h at 37°C in a Thermomixer and then plated. 48 h after transduction, blasticidin (concentrations ranging from 7-15 μg/ml depending on cell lines) was added in the culture media.

### Virus and infections

The SARS-CoV-2 strain BetaCoV/France/IDF0372/2020 (historical) and hCoV-19/France/PDL-IPP01065/2021 (20H/501Y.V2 or SA) were supplied by the French National Reference Centre for Respiratory Viruses hosted by Institut Pasteur (Paris, France) and headed by Pr. S. van der Werf. The human samples from which the historical and South African strains were isolated were provided by Dr. X. Lescure and Pr. Y. Yazdanpanah from the Bichat Hospital, Paris, France and Dr. Vincent Foissaud, HIA Percy, Clamart, France, respectively. These strains were supplied through the European Virus Archive goes Global (EVAg) platform, a project that has received funding from the European Union’s Horizon 2020 research and innovation program under grant agreement #653316. The hCoV-19/Japan/TY7-501/2021 strain (20J/501Y.V3 or Brazil) was kindly provided by Jessica Vanhomwegen (Environment and Infectious Risks Research and Expertise Unit; Institut Pasteur). Viral stocks were produced by amplification on Vero E6 cells, for 72 h in DMEM supplemented with 2% FBS and 1% P/S. The cleared supernatant was stored at −80°C and titrated on Vero E6 cells by using standard plaque assays to measure plaque-forming units per mL (PFU/mL). Cells were infected at the indicated multiplicities of infection (MOI) in DMEM without FBS. Virus inoculum was either removed after 6 h and replaced or topped up with FBS containing culture medium to a final concentration of 2% FBS and 1% P/S. For infections with proteolytically activated SARS-CoV-2, cell monolayers were washed twice with PBS before adding virus inoculum in DMEM supplemented with 1μg/ml of trypsin TPCK (Sigma) and no FBS. After 4h, DEMEM containing FBS was added to a final concentration of 2%.

### TCID_50_ assays

Supernatants of infected cells were 10-fold serially diluted in DMEM supplemented with 2% FBS and 1% P/S. To remove cytokines and other proteins, supernatants were ultracentrifuged for 1 h at 45k rpm at 4°C and resuspended in DMEM with 2% FBS and 1% P/S after 4 h incubation at 4°C. Infected cells were lysed and scraped in ddH_2_O. After one freeze-thaw cycle, whole cell lysates were cleared by centrifugation, supplemented with 10x PBS to a physiological condition and used for serial dilutions. Around 9×10^3^ Vero E6 cells and 50 μl of serially diluted virus suspensions were deposited in 96-well plate in quintuplicate wells. Cells were fixed with 4% paraformaldehyde (PFA) for 30 min at RT and revealed with crystal violet 5 days later. Cytopathic effects (CPE) were assessed by calculating the 50% tissue culture infective dose (TCID50) using the Spearman-Karber method [77].

### Flow cytometry

Cells were detached with trypsin or versene for hACE2 staining. Cells were then fixed in 4% PFA for 30 min at 4°C and staining was performed in PBS, 2% BSA, 2mM EDTA and 0.1% Saponin (FACS buffer). Cells were incubated with goat pAB anti-hACE2-647 (1:100, FAB933R R&D Systems) and/or with antibodies recognizing the spike protein of SARS-CoV (anti-S, 1:1000, GTX632604 Genetex) or anti-S mAb10 (1 μg/ml, a kind gift from Dr. Hugo Mouquet, Institut Pasteur, Paris, France) and subsequently with secondary antibodies anti-human AlexaFluor-647 (1:1000, A21455 Thermo), anti-mouse AlexaFluor-488 (1:1000, A28175 Thermo) or Dylight488 (1:100, SA5-10166 Thermo) for 30 min at 4°C. Cells were acquired on an Attune NxT Flow Cytometer (Thermo Fisher) and data analyzed with FlowJo software v10 (TriStar).

### RNA extraction and RT-qPCR assays

Total RNA was extracted from cells with the NucleoSpin RNA II kit (Macherey-Nagel) according to the manufacturer’s instructions. First-strand complementary DNA (cDNA) synthesis was performed with the RevertAid H Minus M-MuLV Reverse Transcriptase (Thermo Fisher Scientific) using random primers. For batACE2 determination, total RNA was treated with DNAse I (DNAse-free kit, Thermo Fisher Scientific) for 30 min at 37°C before cDNA synthesis with SuperScript IV reverse transcriptase. Quantitative real-time PCR was performed on a real-time PCR system (QuantStudio 6 Flex, Applied Biosystems) with Power SYBR Green RNA-to-CT 1-Step Kit (Thermo Fisher Scientific). Data were analyzed using the 2-ΔΔCT method, with all samples normalized to GAPDH. Genome equivalent concentrations were determined by extrapolation from a standard curve generated from serial dilutions of the pcDNA3.1-hACE2 plasmid (addgene, 145033) or plasmids encoding a fragment of the RNA-dependent RNA polymerase (RdRp)-IP4 of SARS-CoV-2 or a fragment of the ACE2 genome of each bat species. Primers used for RT-qPCR analysis are given in table S1.

### Cloning of qPCR amplicon

To quantify the amounts of bat ACE2 in each cell line, plasmids containing the qPCR amplicon obtained with the primers described in table S1 were generated via TOPO cloning. Briefly, total RNA was extracted from a cadaver of *Myotis myotis* stored at the University of Veterinary and Pharmaceutical Sciences in Brno. For the remaining two bat species, total RNA extracted from NnKi and FLG-R cells were used. RNA was treated for 30 min at 37°C with DNAse I and cDNA synthesized with SuperScript IV reverse transcriptase. These cDNAs were then used as template for PCR amplification of the qPCR bACE2 amplicon using the primers in table S1 and Phusion High-fidelity DNA Polymerase (Thermo). PCR products were gel-purified (NucleoSpin gel and PCR clean-up kit, Macherey-Nagel) and cloned into pCR-Blunt II-TOPO vectors using the Zero Blunt TOPO PCR Cloning Kit (Thermo). Inserts were verified via Sanger sequencing. Plasmids were then used as quantitative qPCR standards.

### Western blot analysis

Proteins extracted from cell lysates were resolved by SDS-polyacrylamide gel electrophoresis on 4-12% NuPAGE Bis-Tris Gel (Life Technologies) with MOPS running buffer and semi-dry transferred to a nitrocellulose membrane with Trans-Blot Turbo system (Bio-Rad). After blocking with 0.05% Tween20 in PBS (PBST) containing 5% dry milk powder for 1 h at room temperature (RT), the membrane was incubated with goat pAB anti-hACE2-700 (1:200, FAB933N R&D Systems) and mouse mAB anti-b-actin (1:5000, A5316 Sigma) diluted in blocking buffer overnight at 4°C. The membranes were then incubated with DyLight800 secondary AB (1:5000, 46421 Thermo) diluted in blocking buffer for 1 h. Finally, the membranes were revealed using an Odyssey CLx infrared imaging system (LI-COR Bioscience).

### Immunofluorescence microscopy and live cell imaging

Cells grown on glass coverslips were fixed in 4% PFA for 30 min at RT and permeabilized with 0.2% Triton X-100 (Sigma/Merck) in PBS for 10 min at RT. Following blocking with 3% bovine serum albumin (BSA, Sigma) in PBS for 1 h at RT, cells were incubated with goat pAB anti-hACE2 (1:50, AF933 R&D Systems) and mAB anti-SARS-CoV-2-spike (1:1000, GTX632604 Genetex) in 1% BSA in PBS (AB buffer) for 1h at RT or overnight at 4°C. Subsequently, cells were incubated with anti-goat Alexa488 (A-11055, Thermo Fisher Scientific) and anti-mouse Alexa555 (A21427, Thermo Fisher Scientific) secondary antibodies diluted 1:500 in AB buffer for 30 min at RT. Finally, cells were stained with NucBlue Fixed Cell ReadyProbes reagent (Thermo) in PBS for 5 min at RT. Coverslips were washed with ultrapure water (Gibco) and mounted in ProLong Gold antifade (Life Technologies). Sample were visualized with a Leica TCS SP8 confocal microscope (Leica Microsystems) and a white light excitation laser and a 405nm diode laser were used for excitation. Confocal images were taken with automatically optimized pixel format, a 4× frame averaging and a scan speed of 400 Hz through an HC PL APO CS2 63x NA 1.4 oil immersion objective. Overlay pictures of single channel images were digitally processed in Leica LAS X lite software. For live imaging, 5.4×10^4^ to 10^5^ cells were plated per quadrant in a μ-Disli 35 mm Quad dish (80416, Ibidi). Cells were infected the next day with SARS-CoV-2 at a MOI of 1 in culture media supplemented with 2.5% FBS and 1% P/S containing propidium iodide. Transmission and fluorescence images were taken at 37°C every 15 min, up to 48 h, using a Nikon BioStation IMQ, with three fields for each condition.

### Attachment and entry assays

Cells plated in monolayers were pre-chilled on ice for 10 min and washed once with cold PBS. Cells were then incubated with SARS-CoV-2 at a MOI of 1 for 1 h on ice. Following three washes with cold PBS, half of the cells was lysed in RA1 lysis buffer (Macherey-Nagel) (“on ice”). The second half of the cells was trypsinized for 15 min on ice and 15min at 37°C after washing of the virus inoculum, then washed with PBS and lysed (“on ice + trypsin”). The remaining cells were directly transferred to 37°C after washing of the virus inoculum and incubated for 2 or 6 h in warm culture media supplemented with 2% FBS and 1% P/S. After this incubation period, those cells were trypsinized for 30 min at 37°C, washed with PBS and lysed in RA1 buffer (“2h”, “6h”). Finally, total RNA was extracted from all cell lysates using the NucleoSpin RNA II kit (Macherey-Nagel).

### Polyl:C stimulation

Cells were plated in monolayers in 24-well culture plates. The next day, they were transfected with 250 ng low molecular weight Poly I:C (InvivoGen) or PBS, respectively, using INTERFERin (Polyplus transfection) transfection reagent. Cells were lysed 16 h after transfection and total RNA was extracted using the NucleoSpin RNA II kit (Macherey-Nagel).

### Statistical analysis

Graphical representation and statistical analyses were performed using GraphPad Prism Version 9.0.2 software (GraphPad). Unless otherwise stated, results are shown as means ± SD from 3 independent experiments. Significance was calculated using either Dunnett’s multiple comparison test on a two-way ANOVA analysis or Šídák’s multiple comparisons test on a two-way ANOVA analysis as indicated. Statistically significant differences are indicated as follows: *p < 0.05; **p < 0.01; ***p < 0.001; ****p < 0.0001; and ns, not significant.

## Acknowledgments

We thank Noémie Aurine (Université Lyon 1, France) for her help designing RT-qPCR primers for bat samples; Bertrand Pain (Université Lyon 1, France) for his advices in both bat primers design and generation of bat primary cell lines; Ondine Filippi-Codaccioni and Marc López-Roig (Université Lyon 1, France) for their precious help in bat sampling; the French National Reference Centre for Respiratory Viruses hosted by Institut Pasteur (France) and headed by Pr. S. van der Werf for providing the historical viral strains and the SA variant; Jessica Vanhomwegen (Institut Pasteur, France) for providing the Brazilian variant; Hugo Mouquet and Cyril Planchais (Institut Pasteur, France) for providing anti-S antibodies; Françoise Porrot for lentiviral production (Institut Pasteur, France); Florence Guivel-Benhassine (Institut Pasteur, France) for help in titration assays; Thomas Vallet (Institut Pasteur, France) for help performing experiments with the viral variants and Matthias Lenk (Friedrich-Loeffler-Institut, Germany) for providing the *E. serotinus* cell lines. We are grateful to the members of our laboratories for helpful discussions. We acknowledge the UTechS Photonic BioImaging (Imagopole), C2RT, Institut Pasteur, supported by the French National Research Agency (France BioImaging; ANR-10-INBS-04; Investments for the Future) for the use of the confocal microscope.

## Funding

This work was funded by the CNRS (NJ, OS), Institut Pasteur (NJ, LD, OS), ‘Urgence COVID-19’ fundraising campaign of Institut Pasteur (NJ, LD, OS), the University of Veterinary Sciences of Brno (FVHE/Pikula/ITA2021) (JP), LabEx Ecofect (ANR-11-LABX-0048) (DoP), Labex IBEID (ANR-10-LABX-62-IBEID) (OS), ANR/FRM Flash Covid PROTEO-SARS-CoV-2 (OS) and IDISCOVR (OS). S.M.A. and De.P. are supported by the Pasteur-Paris University (PPU) International PhD Program and the Vaccine Research Institute (ANR-10-LABX-77), respectively. D.S.L. was funded by the Chinese Scholarship Council and Institut Pasteur. The funders had no role in study design, data collection and analysis, decision to publish, or preparation of the manuscript.

## Author contributions

S.M.A., L.D. and N.J. designed the study. S.M.A, F.S., M.C., De.P. and D.S.L. performed experiments. J.B. generated A549-hACE2 cells and hACE2 lentiviral vectors. J.P., M.N. and V.S. generated the Nn cell lines. J.S.C., Do.P. J.Z. and J.P. organized field trips and collected bat wing biopsies. L.D. generated primary bat cells. S.M.A., F.S., M.C., De.P, L.D. and N.J. analyzed the data. O.S., L.D. and N.J. supervised the work. S.M.A., F.S. and N.J. wrote the manuscript. All authors edited the manuscript.

## Competing interests

The authors declare that no competing interests exist.

## Movie legends

**Movie 1. Time-lapse microscopy of mock-infected MmBr-ACE2 cells**. MmBr-ACE2 cells were seeded at 9×10^4^ cells per quadrant in a μ-Dish 35 mm Quad dish (Ibidi) and cultured in fresh media (2,5% FBS) containing propidium iodide the next day. Transmission and fluorescence images were taken every 15 min, up to 48 h, using a Nikon BioStation IMQ, at 37°C with three fields of acquisition for each condition.

**Movie 2. Time-lapse microscopy of MmBr-ACE2 cells infected with SARS-CoV-2**. MmBr-ACE2 cells were seeded at 9×10^4^ cells per quadrant in a μ-Dish 35 mm Quad dish (Ibidi) and infected the next day with SARS-CoV-2 at a MOI of 1 in culture medium (2,5% FBS) containing propidium iodide. Transmission and fluorescence images were taken every 15 min, up to 48 h, using a Nikon BioStation IMQ, at 37°C with three fields of acquisition for each condition.

**Movie 3. Time-lapse microscopy of mock-infected FLG-ID-ACE2 cells**. FLG-ID cells were seeded at 5.4×10^4^ cells per quadrant in a μ-Dish 35 mm Quad dish (Ibidi) and cultured in fresh media (2,5% FBS) containing propidium iodide the next day. Transmission and fluorescence images were taken every 15 min, up to 48 h, using a Nikon BioStation IMQ, at 37°C with three fields of acquisition for each condition.

**Movie 4. Time-lapse microscopy of FLG-ID-ACE2 cells infected with SARS-CoV-2**. FLG-ID cells were seeded at 5.4×10^4^ cells per quadrant in a μ-Dish 35 mm Quad dish (Ibidi) and infected the next day with SARS-CoV-2 at a MOI of 1 in culture medium (2,5% FBS) containing propidium iodide. Transmission and fluorescence images were taken every 15 min, up to 48 h, using a Nikon BioStation IMQ, at 37°C with three fields of acquisition for each condition.

## Supplementary figure legends

**Fig. S1. Resistance to infection with SARS-CoV-2 variants (B1.351 and P1) in selected bat cell lines. A, B)** Caco-TC7 and bat cell lines were left uninfected (Mock) or were infected with the SARS-CoV-2 variants B1.351 (South Africa) and P1 (Brazil) at a MOI of 1 for 24 hours. The percentages of the indicated cells that express the viral S proteins were determined by flow cytometric analysis. **C)** Caco-TC7 and NnKi cells were left uninfected (Mock) or were infected with SARS-CoV-2 at a MOI of 1 in the absence of FBS and in the presence of trypsin TPCK at 1μg/ml. The percentages of S-positive cells were determined by flow cytometric analysis. Data points represent two technical replicates.

**Fig. S2. Flow cytometry analysis of infected bat cells. A,** Dot plots and density plots of MmBr-ACE2 cells infected with SARS-CoV-2 at a MOI of 0.04 for 24 hours. Cell granularity (SSC) is displayed against size (FSC) to visualize the larger-size subpopulation of cells appearing during infection. **B,** Dot plots of SARS-CoV-2 infected MmBr-ACE2 (MOI of 0.04) and FLG-ID-ACE2 (MOI of 1) cells for 24 hours and stained with anti-spike and anti-hACE2-647 antibodies, displayed on x- and y-axes respectively, to show absence/presence of double positive cell subpopulations. (a,b) Plots are representative of three independent experiments.

## Notes

### Competing Interest Statement

The authors have declared no competing interest.

