## supplementary information for "Species-specific molecular barriers to SARS-CoV-2 replication in bat cells"

### Figure S1

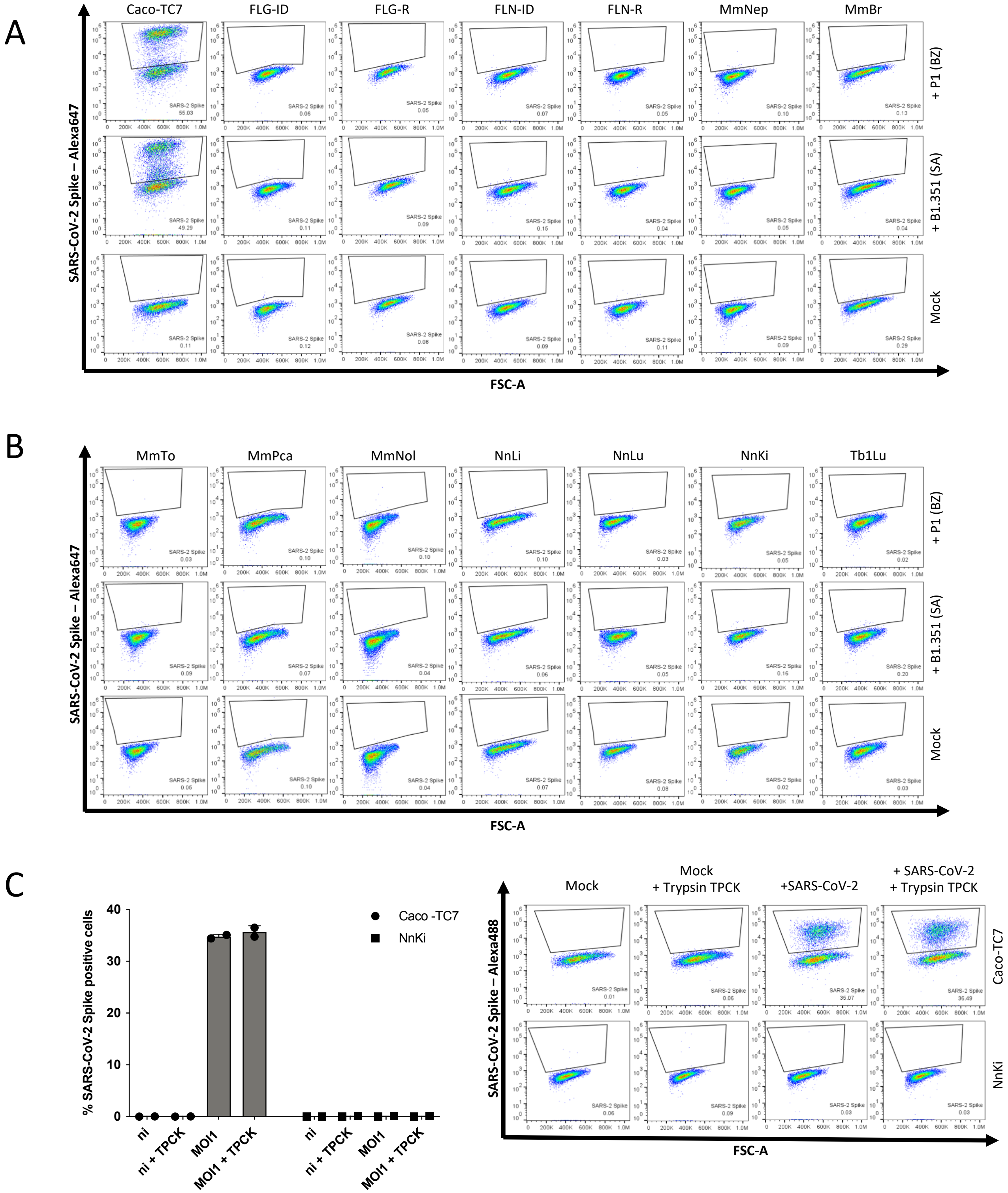
A
Caco-TC7
FLG-ID
FLG-R
FLN-ID
FLN-R
MmNep
MmBr
+ P1 (BZ)
+ B1.351 (SA)
Mock
SARS-CoV-2 Spike – Alexa647
FSC-A
B
MmTo
MmPca
MmNol
NnLi
NnLu
NnKi
Tb1Lu
C
% SARS-CoV-2 Spike positive cells
Caco -TC7
NnKi
ni
ni + TPCK
MOI1
MOI1 + TPCK
Mock
Mock + Trypsin TPCK
+SARS-CoV-2
+ SARS-CoV-2 + Trypsin TPCK
SARS-CoV-2 Spike – Alexa488

### Figure S2

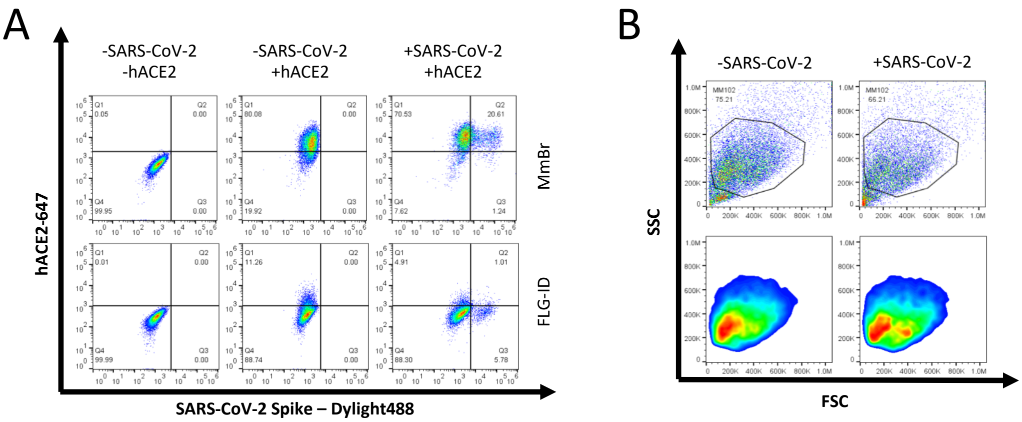


### Table S1. qPCR primers.

| target gene | forward primer | reverse primer |
|---|---|---|
| ACE2 human | GGACCCAGGAAATGTTCAGA | GGCTGCAGAAAGTGACATGA |
| ACE2 FLG | TGGGACTCTACCGTTCACTTA | GCTTCATCTCCCACCACTTT |
| ACE2 Mm | TGCTTATGTCAGGGCAAAGT | CCCACATATCACCAAGCAAATG |
| ACE2 Nn | CAGTCCTGGGATGCAGATAAG | TGGCTCAGTTAGCATGGATTTA |
| GAPDH human | GGTCGGAGTCAACGGATTTG | ACTCCACGACGTACTCAGCG |
| GAPDH FLG | TCATCAACGGAAAGTCCATCTC | ACATACTCAGCACCAGCATC |
| GAPDH Mm | GTAGTGAAGCAGGCATCAGAG | GGAGTGGGTGTCACTGTTAAA |
| GAPDH Nn | CCTGTTCGTCAGACAGCCTT | TTGATGGCGACAACTTGCAC |
| IFIH1 human | ACA CGT TCT TTG CGA TTT CC | ACC AAA TAC AGG AGC CAT GC |
| IFIH1 Mm/FLG | GGAGTCAAAGCCCACCATCT | TCCAGACCTTCTTCTGCCAC |
| IFIH1 Nn | TTTGCCAAGTGAGCCCAATG | AAGCGGTCTTTGCGATTTCC |
| OAS human | GAGCTCCTGACGGTCTATGC | TTCGTGAGCTGCCTTCTCAG |
| OAS pan-bat | GGAAGGAGGGCGAGTTCTC | GGTACCAGTGCTTGACCAGG |
| SARS-CoV-2 nsp12 polymerase | GGTAACTGGTATGATTTC | CTGGTCAAGGTTAATATAGG |
